# Zymocin-like killer toxin gene clusters in the nuclear genomes of filamentous fungi

**DOI:** 10.1101/2024.10.25.620242

**Authors:** Padraic G. Heneghan, Letal I. Salzberg, Kenneth H. Wolfe

## Abstract

Zymocin-like killer toxins are anticodon nucleases secreted by some budding yeast species, which kill competitor yeasts by cleaving tRNA molecules. They are encoded by virus-like elements (VLEs), cytosolic linear DNA molecules that are also called killer plasmids. To date, toxins of this type have been found only in budding yeast species (Saccharomycotina). Here, we show that the nuclear genomes of many filamentous fungi (Pezizomycotina) contain small clusters of genes coding for a zymocin-like ribonuclease (γ-toxin), a chitinase (toxin α/β-subunit), and in some cases an immunity protein. The γ-toxins from *Fusarium oxysporum* and *Colletotrichum siamense* abolished growth when expressed intracellularly in *S. cerevisiae*. Phylogenetic analysis of glycoside hydrolase 18 (GH18) domains shows that the chitinase genes in the gene clusters are members of the previously described C-II subgroup of Pezizomycotina chitinases. We propose that the Pezizomycotina gene clusters originated by integration of a yeast-like VLE into the nuclear genome, but this event must have been ancient because (1) phylogenetically, the Pezizomycotina C-II chitinases and the Saccharomycotina VLE-encoded toxin α/β subunit chitinases are sister clades with neither of them nested inside the other, and (2) many of the Pezizomycotina toxin cluster genes contain introns, whereas VLEs do not. One of the toxin gene clusters in *Fusarium graminearum* is a locus that has previously been shown to be under diversifying selection in North American populations of this plant pathogen. We also show that two genera of agaric mushrooms (Basidiomycota) have acquired toxin gene clusters by horizontal transfers from different Pezizomycotina donors.

## Introduction

Some budding yeast species (Saccharomycotina) contain linear dsDNA molecules in their cytosol that encode killer toxins (Satwika et al., 2012a; Schaffrath et al., 2018). These DNA molecules were originally called killer plasmids, but they have more recently been called virus-like elements (VLEs) to reflect their evolutionary relatedness to eukaryotic viruses with DNA genomes (Satwika et al., 2012a; Krupovic and Koonin, 2015; Koonin et al., 2024). The toxins are secreted by killer cells containing the VLE. The toxins enter victim cells, where they cause cell death by cleaving either tRNA molecules or rRNA molecules. The best-known toxin of this type is zymocin, encoded by a VLE in the yeast *Kluyveromyces lactis*, which cleaves tRNA-Glu(UUC) (Lu et al., 2005; Lu et al., 2008). Zymocin is a heterotrimer consisting of α, β, and γ subunits. The zymocin γ subunit (also called γ-toxin) is the tRNA-cleaving ribonuclease. The α and β subunits are made by proteolytic processing of a single precursor protein which is a chitinase or chitinase-like protein (also called the chitin-binding protein, CBP). Their role is to deliver the γ-toxin to the surface of the victim cell, by binding to chitin in its cell wall (Jablonowski et al., 2001). Both the γ-toxin and the chitinase have N-terminal secretion signals and are secreted by the killer cell, linked by a disulfide bond (Stark et al., 1990). Killer cells avoid being killed by their own toxin because the VLE also encodes an immunity protein that is not secreted (Kast et al., 2015). The immunity protein acts as an antitoxin, neutralizing any toxin molecules that are taken up by the killer cell. In addition to zymocin, three other budding yeast VLE-encoded killer toxins (PaT, DrT and PiT) have been characterized in detail. Zymocin, PaT and DrT cleave tRNAs, whereas PiT cleaves rRNA (Klassen et al., 2008; Chakravarty et al., 2014; Kast et al., 2014).

The chitinases that form α/β subunits of Saccharomycotina killer toxins are modular proteins with multiple domains, typically including a chitin-binding domain, a chitinase (GH18) domain, and a LysM N-acetylglucosamine binding domain (accessions ChtBD1, Glyco_18 and LysM in the SMART database) (Stergiopoulos et al., 2012; Tzelepis and Karlsson, 2019). Chitinases are enzymes that hydrolyze chitin, a polymer of N-acetylglucosamine that is a component of the cell walls of fungi and the exoskeletons of insects and crustaceans (Vega and Kalkum, 2012). Fungal cell walls are polymers of β-(1,3) and β-(1,6) glucans linked to chitin by a β-(1,4) glycosidic bond (Latgé, 2007). Whereas the chitinase domain of the killer toxin α/β subunits is probably used to weaken the victim’s cell wall and enable the γ-toxin to pass through it, other chitinases encoded by the nuclear genome play important roles in other aspects of fungal biology. For example, hydrolysis of cell wall chitin is required during hyphal growth (Seidl, 2008). Chitinases are also virulence factors used by many fungal pathogens to attack competitor fungi or to invade host exoskeletons (Lovett and St. Leger, 2018; Loc et al., 2020; Ma et al., 2024). Some chitinases in pathogenic fungi include an effector domain – a toxic domain that directly harms the host cells, for example the Hce2 domain (Gruber and Seidl-Seiboth, 2012; Stergiopoulos et al., 2012; Tzelepis et al., 2014).

All fungal chitinases contain a GH18 glycoside hydrolase domain (Murphy et al., 2011; Hartl et al., 2012; Tzelepis and Karlsson, 2019). Based on their sequences and domain organization, the fungal GH18 chitinases were divided into three groups (A, B, and C), each with multiple subgroups (Seidl et al., 2005; Karlsson and Stenlid, 2008; Seidl, 2008). More recent analysis based on larger numbers of sequences has indicated that group C, which can be subdivided into C-I and C-II subgroups, is nested within group A (Goughenour et al., 2021). Group C chitinases have structural and sequence similarity to the zymocin chitinase (α/β subunit precursor) and have been termed “killer toxin-like chitinases” (Seidl et al., 2005; Tzelepis et al., 2014; Tzelepis and Karlsson, 2019). Fungi that are soil-borne generally have more chitinase genes than fungi that are human pathogens, and fungi with a mycoparasitic lifestyle have higher numbers of killer toxin-like chitinases, which suggests that some of their multiple chitinases probably have roles in antagonistic fungal-fungal interactions rather than pathogenic interactions against plant or animal hosts (Karlsson and Stenlid, 2008; Tzelepis and Karlsson, 2019).

In a recent study, we used bioinformatic methods to search systematically for new zymocin-like killer toxin genes in genome sequence data from budding yeasts (Saccharomycotina) and found 45 candidates, many of which were confirmed as functional (Heneghan et al., 2024). Some of these new toxins are encoded on VLEs like in *K. lactis*, and others are in nuclear genomes. In most cases, whether located on a VLE or in the nuclear genome, the toxin gene is beside genes for a chitinase and an immunity protein. During these searches, we noticed that some of the Saccharomycotina immunity genes have homologs in Pezizomycotina genomes, which motivated us to investigate the Pezizomycotina loci in more detail. We show here that many Pezizomycotina species have nuclear gene clusters encoding a zymocin-like toxin candidate, a chitinase, and a putative immunity protein, and that some species have multiple clusters coding for diverse toxins. The chitinases correspond to the previously described killer toxin-like C-II subgroup. Phylogenetically the C-II chitinases form a sister clade to the chitinases encoded by Saccharomycotina VLEs, which indicates that they have a shared evolutionary origin. Our work verifies that C-II chitinases truly are killer-like chitinases and strongly suggests that they act in concert with zymocin-like anticodon nucleases, in situations where one fungal cell attacks another. It adds these ribonucleases to the arsenal of molecules known to be used by filamentous fungi for interspecies and inter-individual warfare (Lacadena et al., 2007; Jones et al., 2021).

## Results

### Clusters of immunity, chitinase and putative killer toxin genes in Pezizomycotina genomes

By using TBLASTN searches against NCBI genome sequence databases, we first discovered that immunity (*Imm*) genes from Saccharomycotina VLEs have homologs in some Pezizomycotina genomes such as *Colletotrichum siamense* (taxonomic class Sordariomycetes) and *Penicillium robsamsonii* (Eurotiomycetes). These genes are located close to genes for secreted chitinases with GH18 glycoside hydrolase domains, which is reminiscent of the organization of Saccharomycotina VLEs such as the one encoding *Kluyveromyces lactis* zymocin (Figure 1). We then found that the regions beside the chitinase and putative immunity genes contained genes potentially coding for small secreted proteins, so we considered these to be candidate zymocin-like toxins, described in more detail in the next sections. Additional TBLASTN searches using the Pezizomycotina chitinase, candidate toxin, and candidate immunity proteins as queries led to the discovery that many other Pezizomycotina genomes contain clustered chitinase and toxin candidate genes, often without a candidate immunity gene (Figure 1; Table S1). In all but two of the species that lack *Imm* genes in their clusters, there at least one *Imm* gene somewhere in the genome (Table S2).

**Figure 1.**
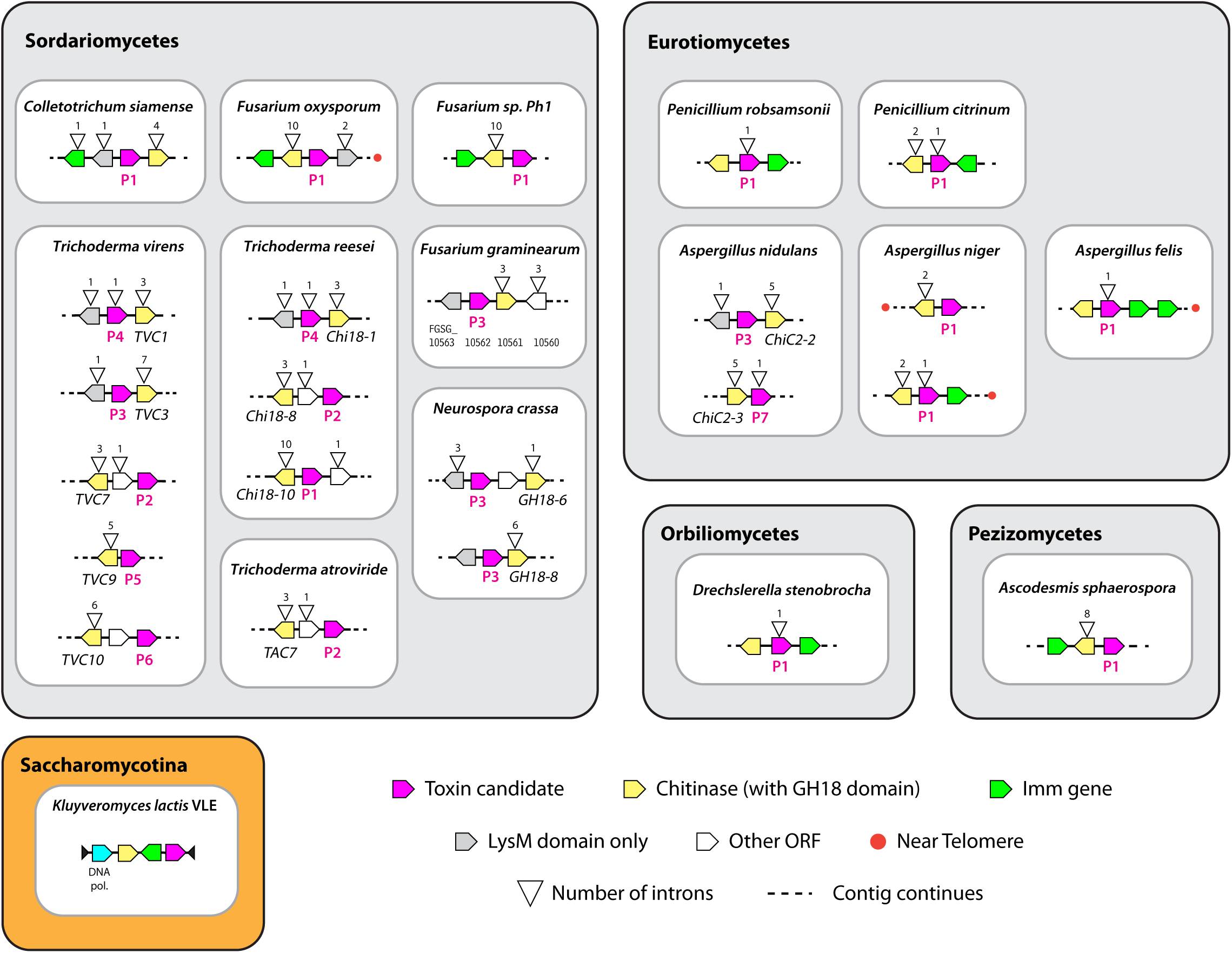
Zymocin-like killer toxin gene clusters in Pezizomycotina species. Examples are shown of some toxin clusters discovered in Sordariomycetes and Eurotiomycetes, which are the two largest taxonomic classes in subphylum Pezizomycotina, as well as all clusters discovered in other classes. The structure of the VLE (cytosolic linear plasmid pGKL1) encoding zymocin in the budding yeast *Kluyveromyces lactis* is included for comparison. Names are shown for chitinase genes whose function has been investigated in previous studies (Seidl et al., 2005; Gruber et al., 2010; Tzelepis et al., 2012; Tzelepis et al., 2014). Labels P1 to P9 indicate the sequence group (Tox-P1 to Tox-P9) of each candidate toxin. Genes containing only LysM (CBM50) domains are marked and appear to belong to the clusters (Gruber et al., 2010). Genes with introns are marked by a triangle showing the number of introns. Red dots indicate clusters that are potentially subtelomeric or in repetitive regions within 50 kb of the end of a contig. Not drawn to scale. All clusters are shown in the orientation in which the toxin gene is transcribed from left to right. More information about cluster locations and sequence accession numbers is given in Table S1.

Most of the gene clusters were found in Eurotiomycetes and Sordariomycetes, which are two of the seven taxonomic classes within subphylum Pezizomycotina. Only two clusters were found in other classes: one in *Drechslerella stenobrocha* (Orbiliomycetes), and one in *Ascodesmis sphaerospora* (Pezizomycetes) (Figure 1). We did not find any clusters in Leotiomycetes or Dothideomycetes, both of which have been sequenced extensively (Shen et al., 2020).

Many of the genes in the Pezizomycotina clusters have introns (Figure 1), whereas VLEs do not contain any introns, which indicates that the clusters have been present in Pezizomycotina nuclear genomes for a considerable amount of time; they are not recently-integrated VLE DNA. Interestingly, some well-studied and closely related chitinases like *Trichoderma atroviride TAC7*, *Trichoderma reesei Chi18-8*, and *Trichoderma virens TVC7* (Gruber et al., 2010) occur in clusters with identical organization, with the same introns and gene content, but at different (non-syntenic) genomic locations. Similarly, the gene clusters found in *Fusarium* species were all quite closely related but are not syntenic. The *Colletotrichum siamense* toxin gene cluster (Figure 1) is present at the same location in only a few species in this genus so it appears to have been inserted there relatively recently (Figure S1), whereas the cluster beside *Trichoderma reesei Chi18-10* is at a location conserved across many *Trichoderma* species but has been lost in one subclade of the genus (Figure S2). In summary, toxin gene clusters appear to be easily gained, lost, or moved around the genome.

All but three of the Pezizomycotina chitinases in the clusters have LysM (CBM50) domains (Table S1), similar to the modular structure of Saccharomycotina killer toxin chitinases. In addition to chitinase (GH18), putative toxin, and putative immunity genes, some clusters contain genes coding for proteins with only LysM domains (Figure 1). LysM-only proteins have been shown to be important virulence and avirulence factors in some plant and insect pathogenic fungi (Laugé et al., 1997; Stergiopoulos et al., 2010; Marshall et al., 2011; Akcapinar et al., 2015; Cen et al., 2017). Some Saccharomycotina VLEs also encode LysM-only proteins, such as *Debaryomyces robertsiae* pWR1A ORF4 and *Millerozyma acaciae* pPac1-2 ORF3 (Klassen et al., 2004). We also found two loci at which a putative toxin gene is located beside a different type of hydrolase gene, with no chitinase (GH18 domain) gene nearby: one in *Aspergillus flavus* with a GH71 (mutanase) domain, and one in *Xylaria arbuscula* with a xylanase (XynB-like) domain (Table S1).

### Pezizomycotina killer toxin candidates are functional in *S. cerevisiae*

We tested whether three Pezizomycotina toxin candidates are functional toxins: *Colletotrichum siamense* (strain SAR 10_71; (Rehner et al., 2023)), *Fusarium oxysporum* (*f. sp. pisi*-37622 HDV247, gene FOVG_08799; (Williams et al., 2016)), and *Penicillium robsamsonii* (strain IBT 29466, gene N7447_004436; (Petersen et al., 2023)) (Figure 2). Each gene was synthesized, without its secretion signal, and integrated into the *S. cerevisiae* genome under the control of either a β-estradiol inducible promoter or a galactose-inducible promoter, similar to the assays that have been used previously to test killer toxins encoded by Saccharomycotina VLEs (Klassen et al., 2004; Heneghan et al., 2024). In these assays, functional toxins will inhibit or abolish growth of *S. cerevisiae* when induced, because the toxin is not secreted and no immunity protein is present. We first used a β-estradiol induction system to test two toxin candidates and found that the *C. siamense* toxin candidate inhibited growth of *S. cerevisiae* (Fig. 2A), whereas the *P. robsamsonii* candidate did not (Fig. 2B). We then used a galactose induction system, which appears to be more robust, to re-rest the *C. siamense* candidate and to test the *F. oxysporum* candidate. Both of these proteins abolished growth of *S. cerevisiae* upon induction with galactose (Fig. 2C,D). We conclude that the *C. siamense* and *F. oxysporum* proteins are toxins that are functional against *S. cerevisiae*, whereas the *P. robsamsonii* candidate is nonfunctional.

**Figure 2.**
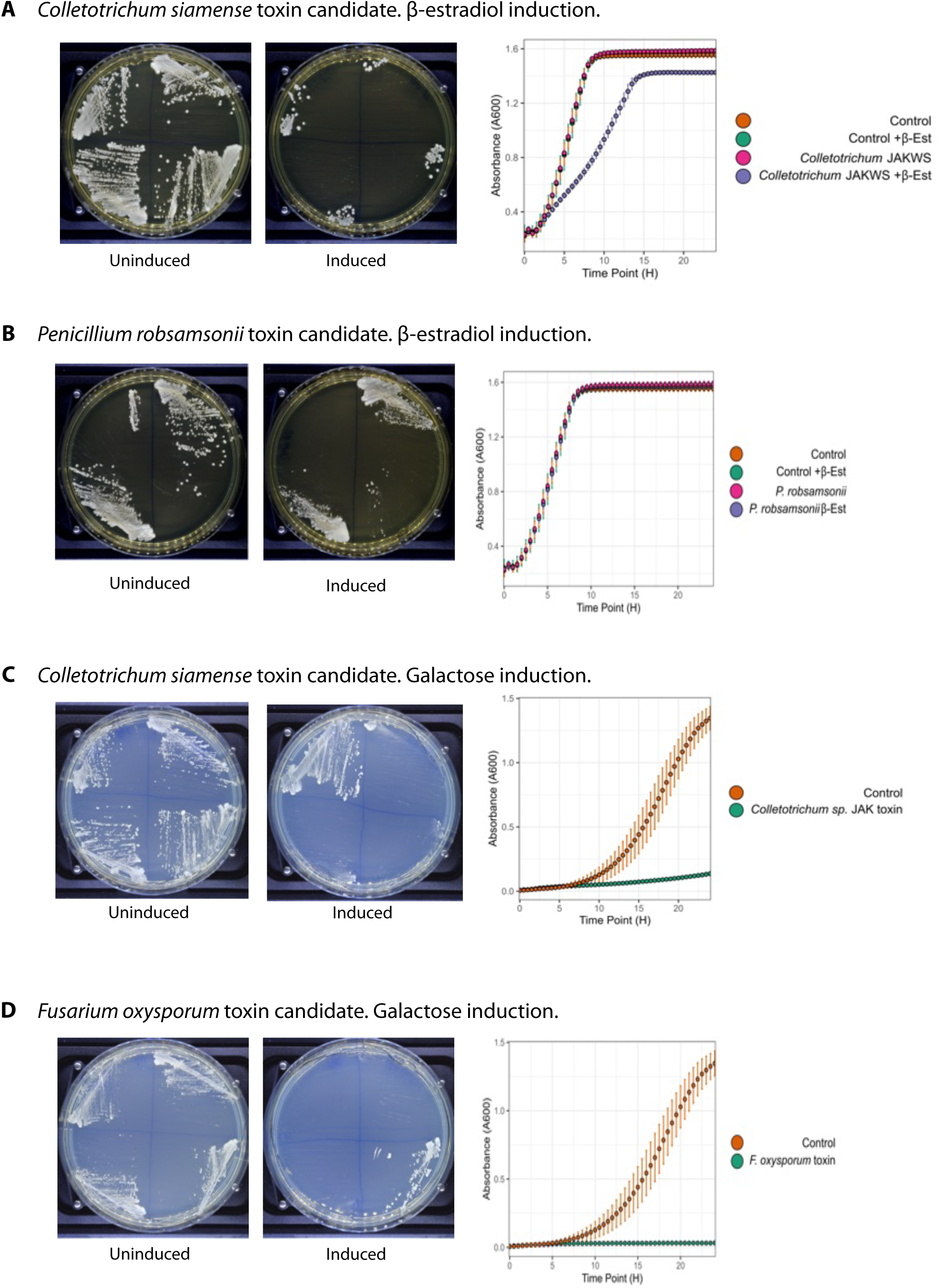
Assays of Pezizomycotina toxin candidate gene function in *S. cerevisiae*. The toxin candidates from *Colletotrichum siamense* and *Fusarium oxysporum* inhibit growth when expressed intracellularly in *S. cerevisiae*, whereas the toxin candidate from *Penicillium robsamsonii* does not. Toxin genes, without their secretion signals and codon-optimized for *S. cerevisiae*, were expressed in *S. cerevisiae* under promoters that are inducible by either β-estradiol (*LexO-CYC1* promoter) or galactose (*GAL1* promoter). For β-estradiol induction (A, B), plate images show 2-4 independent clones of each toxin gene, streaked onto YPD agar (left plates) or YPD + 2 μM β-estradiol (right plates), and graphs show growth curves over 24 h in liquid YPD media ± 2 μM β-estradiol, compared to a control (*S. cerevisiae* strain PHY040) that contains the β-estradiol induction system but has no gene inserted downstream of the inducible promoter. For galactose induction (C, D), plate images show growth of 4 independent clones on SC + 2% raffinose (left) and SC + 2% galactose (right), and graphs show growth curves over 24 h in SC + 2% galactose for toxin-expressing strains compared to a control (*S. cerevisiae* strain LS339). One of the four clones of the *Colletotrichum siamense* toxin was non-functional on galactose agar plates (C) and was not used in the liquid assays.

### Pezizomycotina toxins are very divergent from other zymocin-like toxins

There is no statistically significant sequence similarity between any of the Pezizomycotina toxin candidates and any of the known zymocin-like killer toxins from Saccharomycotina VLEs. This is not surprising because, even within Saccharomycotina, the toxins are highly diverse (Heneghan et al., 2024). Using a liberal BLASTP cutoff of *E* < 1e-6 to construct networks of related sequences that we visualized with Cytoscape, we found that most of the 50 Pezizomycotina toxin candidates we examined fall into a single large sequence group that we named Tox-P1 (Figure 3). We also identified eight smaller Pezizomycotina toxin sequence groups that we named Tox-P2 to Tox-P9. The nine groups are all unrelated to each other at the similarity cutoff used (BLASTP *E* < 1e-6), and unrelated to any of the 12 Saccharomycotina sequence groups we identified previously (Figure 3). It is striking that several species have multiple toxin gene clusters, each coding for a different toxin; for example, the five clusters in *Trichoderma virens* code for toxins in the Tox-P2, -P3, -P4, -P5 and -P6 sequence groups (Figure 1).

**Figure 3.**
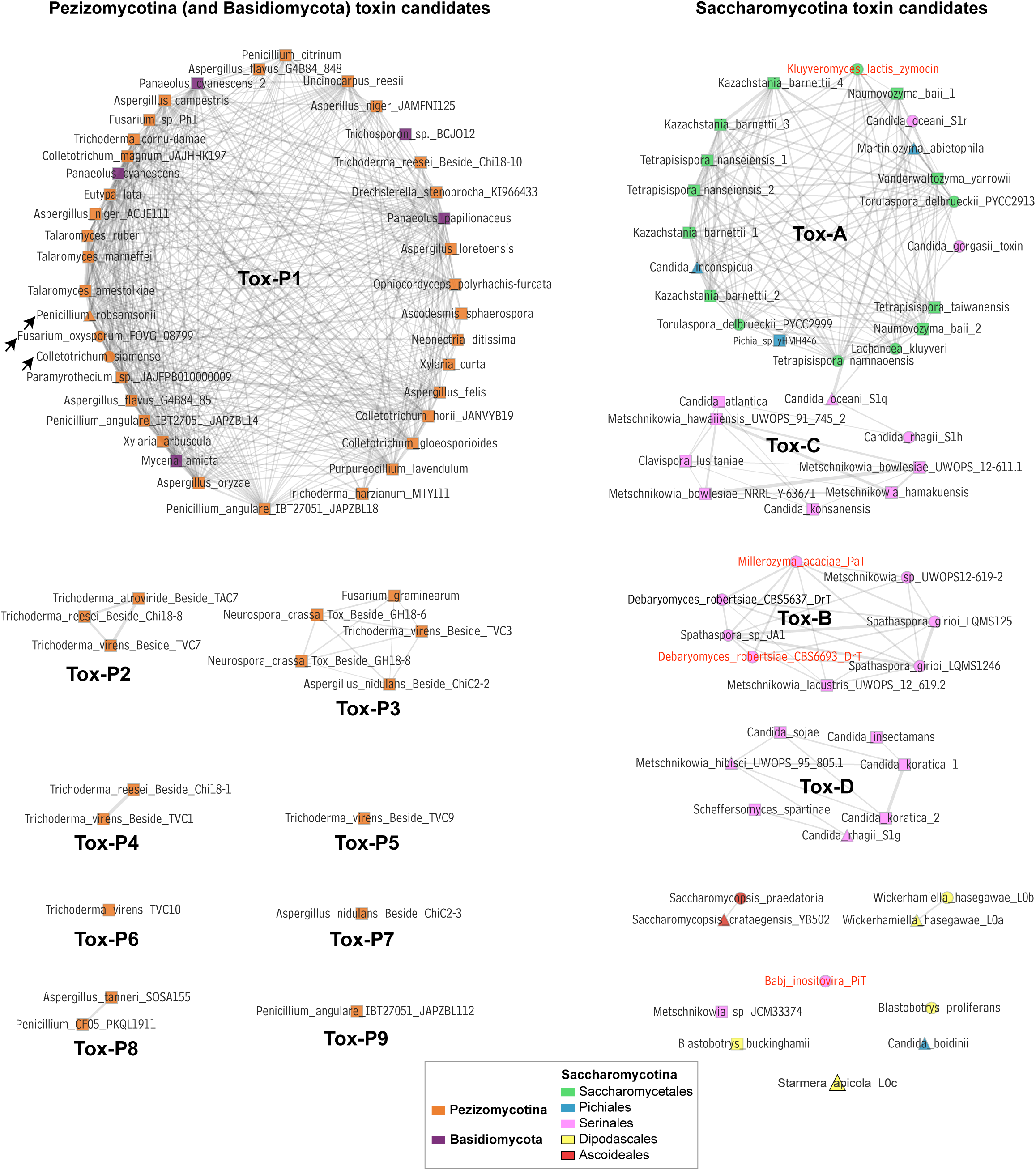
Cytoscape visualization of candidate toxin protein sequence groups. All-vs-all BLASTP searches were performed on candidate toxin proteins to understand their relationship to one another, and the networks were visualized using Cytoscape. Each node represents a candidate toxin, labeled with species and strain or gene information. Node colors indicate taxonomy of the source species, as shown in the key. Two nodes are connected by an edge if the BLASTP hit between that pair of proteins has an *E-*value of 1e-6 or lower; thicker edges indicate lower *E-*values (stronger hits). At this BLASTP threshold, there are no hits between any Pezizomycotina toxins and Saccharomycotina toxins. Most of the Pezizomycotina toxins form a large group (Tox-P1) separate from the Saccharomycotina toxins. Arrows indicate the three toxin candidates in this group that we tested for activity in *S. cerevisiae*. Other Pezizomycotina candidate toxin sequence groups are labeled Tox-P2 to Tox-P9; there is no significant sequence similarity among any of these groups, or to the Tox-P1 group (i.e., all BLASTP scores are *E* > 1e-6) . The Saccharomycotina toxin groups Tox-A, -B, -C and -D identified in our previous study (Heneghan et al., 2024) are marked. Nodes with red labels are the 4 canonical killer plasmid toxins that have been studied in Saccharomycotina (zymocin, PaT, DrT and PiT). Node shapes indicate the results of assays in *S. cerevisiae*: functional toxins (triangles), candidates that were tested but found to be non-functional (circles), and candidates that were not tested (squares).

In Saccharomycotina zymocin-like toxins, a small number of amino acid residues are essential for their ribonuclease activity (Meineke et al., 2012; Chakravarty et al., 2014). One of the most conserved residues is glutamate at the eighth position (Glu-8) of the mature γ-toxin after the secretion signal sequence is cleaved off (Meineke et al., 2012; Chakravarty et al., 2014). All but four of the 50 Pezizomycotina toxin candidates have a Glu-8 residue in the mature protein, when their secretion signal cleavage sites are predicted using TargetP (Emanuelsson et al., 2007) (Table S3).

Additionally, a cysteine residue close to the C-terminus of the toxin protein (γ-subunit) is essential for its disulfide bond to the β-subunit, which is made by proteolytic cleavage of the chitinase (precursor of the α and β subunits) (Wemhoff et al., 2014). All but five Pezizomycotina toxin candidates have a cysteine (Table S3). Similarly, the other partner in this disulfide bond – a cysteine near the C-terminus of the chitinase – is also well conserved in the Pezizomycotina chitinase genes in the clusters. Of the 39 Pezizomycotina chitinases containing GH18 domains in the clusters we examined, only eight lack a terminal cysteine, and five of these eight also lack a neighboring putative γ-toxin gene (Figure S3, Table S1). Of the 31 chitinase genes beside a γ-toxin gene, 29 have a terminal cysteine. The conservation of the cysteines suggests that the Pezizomycotina toxins maintain the same disulfide bond between their β-and γ-subunits as in Saccharomycotina.

### The gene cluster chitinases are in subgroup C-II and form a sister clade to Saccharomycotina VLE chitinases

Using the GH18 domain sequences, we constructed a maximum-likelihood phylogenetic tree to relate the Pezizomycotina chitinases in the toxin gene clusters to other established fungal chitinases (from InterPro’s ‘reviewed’ dataset) and to Saccharomycotina chitinases. The tree (Figure 4) has a similar topology to what has been described previously, resolving the Pezizomycotina chitinases into groups A, B, C-I and C-II (Seidl et al., 2005; Karlsson and Stenlid, 2008; Seidl, 2008). We included the nuclear-encoded chitinases from the yeasts *Saccharomyces cerevisiae* (*CTS1, CTS2*) and *Candida albicans* (*CHT1-CHT4*), and these fall into groups A and B.

**Figure 4.**
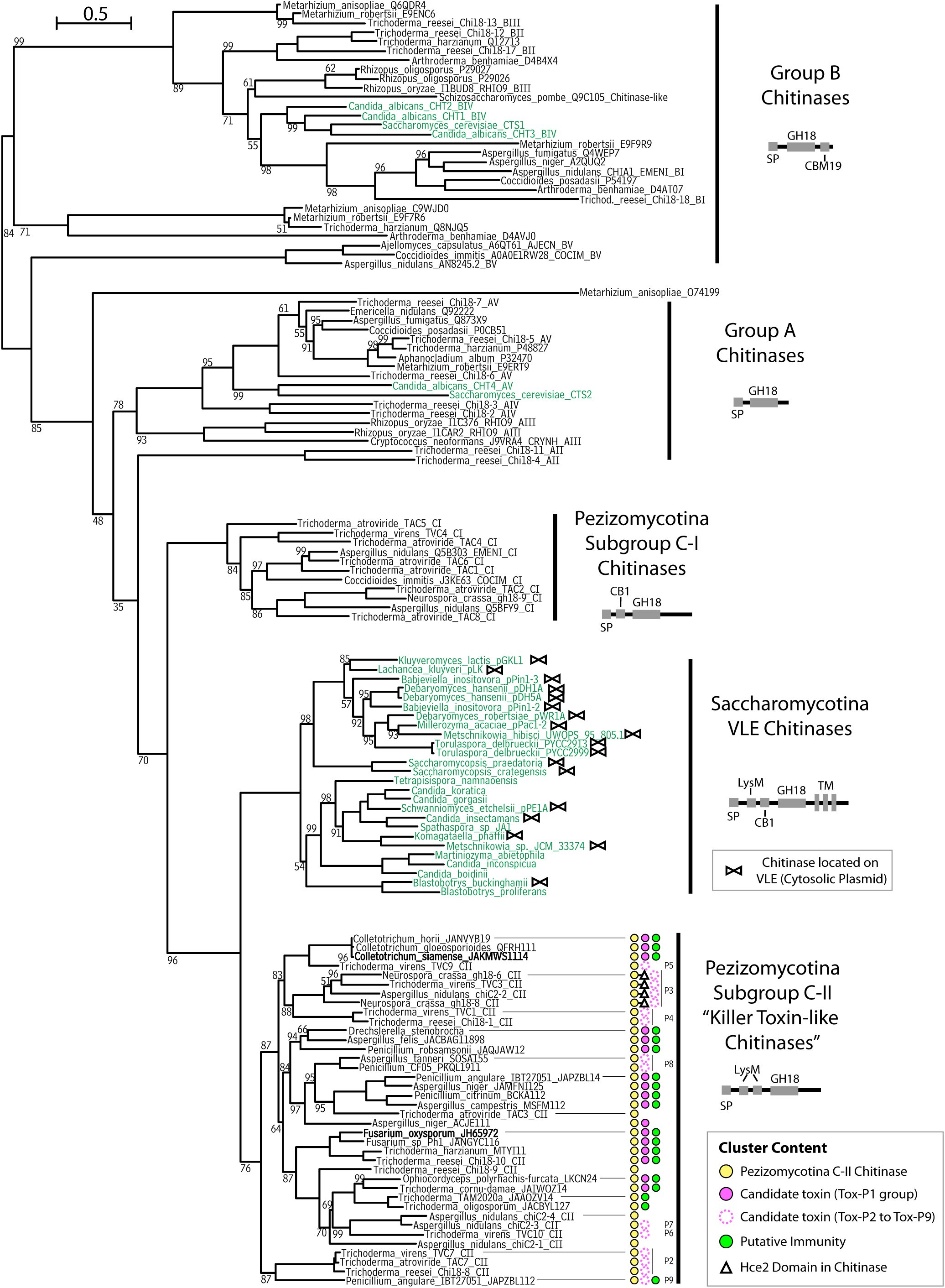
The chitinase genes in Pezizomycotina killer toxin gene clusters are subgroup C-II chitinases. The phylogenetic tree was constructed from an alignment of chitinase GH18 domains from Pezizomycotina and Saccharomycotina (species named in black and green, respectively). It shows that Pezizomycotina subgroup C-II chitinases are a sister clade to the Saccharomycotina chitinases encoded by VLEs or VLE-derived DNA in the nuclear genome. Colored symbols in the Pezizomycotina C-II clade indicate the gene content of the chitinase/toxin/immunity gene clusters present at these loci. Bold species names mark the C-II chitinase genes from *Colletotrichum siamense* and *Fusarium oxysporum* that lie beside the toxin genes we found to be functional against *S. cerevisiae*. Cartoons below each group label illustrate the protein domain structure of a typical member of the group, with domains of interest labeled: SP, signal peptide; GH18, glycoside hydrolase 18 domain; LysM, CBM50 carbohydrate-binding domain (PF01476); CB1, chitin-binding module (PF03427); CBM19, carbohydrate-binding domain (PF00187); TM, transmembrane domain. Branches in the phylogenetic tree have 100% bootstrap support except where shown.

The Pezizomycotina GH18 domains from the toxin cluster chitinases are all members of the previously described C-II subgroup. These Pezizomycotina C-II chitinases form a sister clade to the Saccharomycotina VLE chitinases, with strong bootstrap support (96%; Figure 4). This relationship is also supported by the similarity of domain structures: the C-II and VLE chitinases contain LysM domains, whereas the C-I chitinases do not (Figure 4; (Tzelepis and Karlsson, 2019)). The Saccharomycotina VLE chitinases in this phylogenetic group include some that are located on VLEs (killer plasmids or cryptic plasmids), and others that are located on VLE-derived DNA that recently integrated into yeast nuclear genomes (Heneghan et al., 2024). The phylogenetic sister relationship between Pezizomycotina subgroup C-II chitinases and Saccharomycotina VLE chitinases is consistent with the presence of genes for anticodon nuclease toxins and antitoxin immunity factors beside the chitinase genes in both of these fungal subphyla. However, not all Pezizomycotina C-II chitinase genes are located in toxin gene clusters; we did not find any candidate toxin or immunity genes beside *Trichoderma atroviride TAC3*, *Trichoderma reesei Chi18-9*, and *Aspergillus nidulans chiC2-1* and *chiC2-4* (Figure 4).

### Toxin group Tox-P3 is associated with chitinases containing Hce2 effector domains and is under diversifying selection

The gene content of the gene clusters beside C-II chitinase genes is summarized in Figure 4. Assuming that the history of the toxin genes is the same as the history of their neighboring chitinase genes, the topology of the chitinase phylogenetic tree suggests that Tox-P1, which is the most common toxin sequence group, is ancestral (Figure 4). Each of the other toxin sequence groups Tox-P2 to Tox-P9 is monophyletic, with all of them except Tox-P2 and Tox-P9 being nested within the distribution of Tox-P1.

The chitinases *Aspergillus nidulans ChiC2-2*, *Trichoderma virens TVC3* and *Neurospora crassa GH18-6* and *GH18-8* are unique in that they contain an Hce2 effector domain (Figure 4), whereas none of the other chitinases have this domain. Hce2 domains are homologs of *Cladosporium fulvum* Ecp2, which is a virulence factor in fungal pathogens of plants (Stergiopoulos et al., 2012). The chitinase genes with an Hce2 domain form a monophyletic group, and they are exclusively adjacent to genes for putative killer toxins in the Tox-P3 sequence group. All of these clusters also encode LysM-only effector proteins.

When reviewing the literature on Hce2 domains, we found a report by Kelly and Ward (2018) that a small (10-kb) genomic region that is under strong selection for sequence diversity in North American populations of the plant pathogen *Fusarium graminearum* contains a gene for a C-II chitinase with an Hce2 domain. The region was one of 14 strongly selected regions identified in their study, which used a sliding window approach with non-overlapping 10-kb genomic windows. The identified 10-kb window contains three genes: *FGRRES_10561* (C-II chitinase with Hce2 domain), *FGRRES_10562* (which we find is a putative toxin in the Tox-P3 sequence group) and *FGRRES_10560* (unknown function), and there is also a LysM-only gene (*FGRRES_10563*) just outside the window (Figure 1). Kelly and Ward (2018) highlighted that, when compared among strains of *F. graminearum* and related *Fusarium* species, the three genes in the window stand out as ‘outliers’ whose phylogeny is discordant with the genome-wide SNP phylogeny. For both the toxin and the chitinase there is evidence of trans-species polymorphism: isolates from one North American *F. graminearum* population (population NA3) contain alleles that have introgressed from a different species, *Fusarium gerlachii*. Introgressed alleles are not present in the other two North American populations, resulting in high sequence divergence between them and NA3 at this locus (Kelly and Ward, 2018). Therefore, selection for diversity is acting on the putative toxin gene as well as on the chitinase gene. It is possible that the chitinase in this cluster serves two purposes – zymocin-like killing of competitor fungi, and Hce2-mediated damage to plant host cells.

### Diversity of putative immunity genes

The putative immunity genes we found in Pezizomycotina were identified because they have weak but significant sequence similarity (BLASTP scores in the range *E* = 1e-5 to 1e-10) to the known immunity genes of Saccharomycotina VLEs (*K. lactis* pGKL1 ORF3, *M. acaciae* pPac1-2 ORF4, and *D. robertsiae* pWR1A ORF5) (Chakravarty et al., 2014; Klassen et al., 2014) or their homologs in other Saccharomycotina. When a sequence similarity network is constructed using a more stringent BLASTP cutoff (*E* < 1e-13), the Pezizomycotina genes fall into a single large sequence group, separate from the previously identified Saccharomycotina immunity sequence groups (Heneghan et al., 2024) (Figure S4; Table S4).

All of these Pezizomycotina putative immunity genes are located in gene clusters whose toxin genes are in the Tox-P1 sequence group, except for one that is in a Tox-P9 cluster, and two that are in clusters that do not contain any toxin candidate (Figure 4). This observation suggests that the putative immunity genes we identified probably confer immunity only to Tox-P1 and Tox-P9, and that the genes responsible for immunity to the other classes of toxin (Tox-P2 to Tox-P8) remain unidentified. In some of the gene clusters in the Tox-P2, Tox-P3 and Tox-P6 sequence groups, there are unidentified ORFs located within the gene cluster (Figure 1), making them strong candidates to be immunity genes for these other toxin groups.

### Horizontal transfers of Pezizomycotina toxin gene clusters into basidiomycetes

Most Basidiomycota species do not contain zymocin-like killer toxin gene clusters, but we found four species that do (Figure 5A). These clusters are in the agaric mushroom genera *Panaeolus* and *Mycena* (order Agaricales) and in the basidiomycete yeast *Trichosporon gracile* (order Trichosporonales) (Table S5; Table S6). Their putative toxin genes are all in the Tox-P1 sequence group (Figure 3). *Panaeolus cyanescens* has two clusters, including two toxin genes, arranged as a dispersed direct repeat 24 kb apart. *Panaeolus papilionaceus* also has a dispersed direct repeat, but in this case the putative immunity gene is the only completely duplicated gene (Figure 5A). We identified putative immunity genes only in the two *Panaeolus* species, and these genes are related to the ones in Pezizomycotina (Figure S4). Other genes in the contigs containing each of the four toxin gene clusters are unambiguously of basidiomycete origin, so we can rule out the possibility that the source of the sequenced DNA was misidentified.

**Figure 5:**
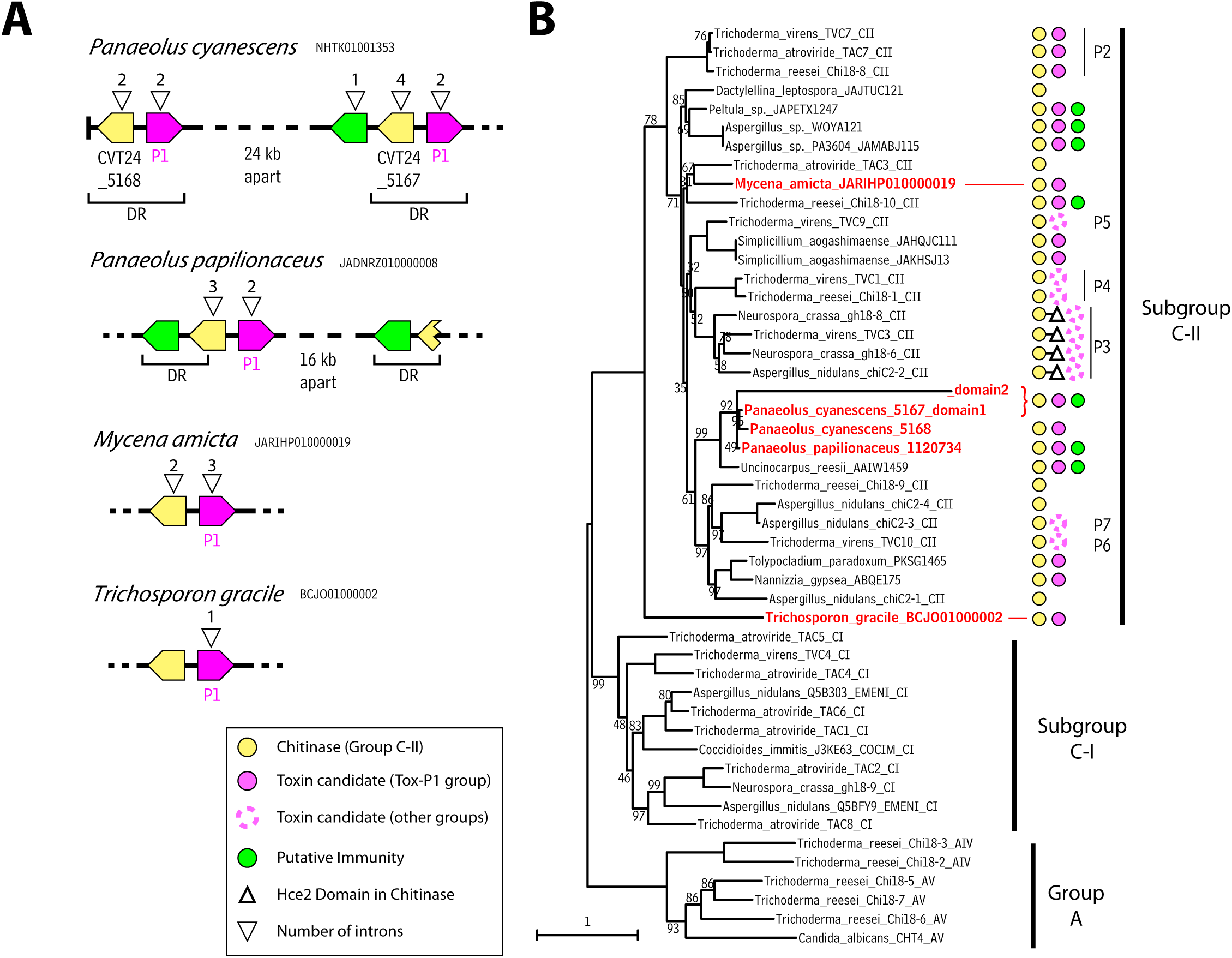
Horizontal transfers of toxin gene clusters from Pezizomycotina to Basidiomycota. **(A)** Toxin gene clusters identified in Basidiomycota. NCBI accession numbers of sequences are shown. DR indicates direct repeats in the *Panaeolus* sequences. **(B)** Phylogenetic tree of GH18 domains, showing the relationship of gene cluster chitinases from Basidiomycota (names in red) to those in Pezizomycotina (names in black). Colored symbols in the C-II clade indicate the gene content of the chitinase/toxin/immunity gene clusters present at these loci. The C-II chitinase sequences in the tree include the strongest matches identified in TBLASTN searches against the NCBI genome sequence databases, using the Basidiomycota chitinases as queries, as well as most of the C-II sequences from Figure 4. The tree was rooted using Group A chitinases. Bootstrap support values are shown for branches having <100% support. Further details of the Basidiomycota clusters are given in Table S5 and Table S6.

The basidiomycete chitinases in the clusters are closely related to Pezizomycotina C-II chitinases, so we constructed a phylogenetic tree using their GH18 domains (Figure 5B). One of the basidiomycete chitinases, *CVT24_005167* in *P. cyanescens*, has two separate GH18 domains, one of which is highly divergent in sequence. In the tree, all four GH18 domains from *Panaeolus* species form a monophyletic group, nested within the Pezizomycotina C-II chitinases (Figure 5B), suggesting that they originated in a single horizontal gene transfer event from a Pezizomycotina donor to the common ancestor of *P. cyanescens* and *P. papilionaceus.* The GH18 domain from the *Mycena amicta* chitinase is also nested within the Pezizomycotina C-II sequences but at a different position, indicating that it has a different Pezizomycotina donor, so we infer that there have been two separate events of horizontal transfer of clusters from Pezizomycotina to Basidiomycota. We did not identify any Pezizomycotina sequences highly similar to the *Panaeolus* or *Mycena* clusters, so there is no “smoking gun” identifying their donors, and in the case of *Panaeolus* the transfer event must have been millions of years ago because it pre-dates the split between *P. cyanescens* and *P. papilionaceus*. The GH18 domain from the *Trichosporon gracile* chitinase is not nested within the Pezizomycotina C-II chitinases in the tree but instead forms a sister lineage to them (Figure 5B). It is therefore unclear whether the *Trichosporon* cluster originated by horizontal transfer.

## Discussion

We have found the first evidence of zymocin-like killer toxins in the fungal subphylum Pezizomycotina. These toxins are diverse with relatively little amino acid conservation, similar to the toxins we found previously in Saccharomycotina (Heneghan et al., 2024). Two of the three Pezizomycotina toxin candidates that we tested showed toxic effects on *S. cerevisiae* when induced. It is unclear why the toxin candidate from *P. robsamsonii* had no effect on *S. cerevisiae*. This protein retains two of the residues known to be essential for ribonuclease activity as described above, and the putative immunity gene in the *P. robsamsonii* gene cluster is related to those neighboring the functional toxin genes in *Colletotrichum siamense* and *Fusarium oxysporum.* It is possible that the *P. robsamsonii* toxin is functional, but that *S. cerevisiae* is not an optimal target for it.

The most probable natural function of the Pezizomycotina zymocin-like toxins is in antagonistic fungal-fungal interactions, because we can infer that the toxins are delivered to recipient cells with chitin in their cell walls. This function directly corroborates previous hypotheses that the C-II lineage of chitinases in Pezizomycotina is involved in fungal-fungal interactions, due to their clear homology to chitinases from Saccharomycotina killer plasmid VLEs (Seidl et al., 2005; Karlsson and Stenlid, 2008; Stergiopoulos et al., 2012; Tzelepis and Karlsson, 2019). It has previously been shown that expression of many group C chitinases, including ones in both the C-I and C-II subgroups, are induced in the presence of other fungi (reviewed by Tzelepis and Karlsson (2019)).

Moreover, *Aspergillus nidulans* shows a reduction in its killing of other fungi when the C-II chitinase *ChiC2-2* is deleted, although it was suggested that this activity was due to the Hce2 domain (Tzelepis et al., 2014). The LysM-only proteins in these clusters can also be coexpressed with the chitinases when a fungus such as *T. atroviride* interacts with fungal prey (Gruber et al., 2010; Akcapinar et al., 2015). If our hypothesis that zymocin-like toxins are used by filamentous fungi to suppress other fungi is correct, these anticodon nucleases may have applications as biocontrol agents for plant protection as previously suggested for the chitinases (Seidl, 2008).

The patchy phylogenetic distribution of zymocin-like toxin gene clusters in Pezizomycotina, their specific association with the C-II subgroup of chitinases, and their sister relationship to the genes on Saccharomycotina killer plasmids (VLEs), beg many questions about the evolutionary history of these genes and clusters. Their origin is like a chicken-and-egg question: (1) were chromosomally encoded gene clusters the ancestors of the killer plasmids, or (2) were killer plasmids the ancestors of the chromosomal clusters? In the first scenario, a piece of chromosomal DNA needs to be converted into an independent cytosolic linear plasmid, which entails gaining the services of a helper plasmid that provides housekeeping functions such as cytosolic DNA polymerase and RNA polymerase (Satwika et al., 2012b). Helper plasmids of this type (such as *K. lactis* pGKL2) have never been found in Pezizomycotina. It also entails losing all introns and gaining protein-capped terminal inverted repeats to form a linear plasmid. Alternatively, in the second scenario, a cytosolic linear killer plasmid needs to become integrated and functional in the nuclear genome, with three genes (toxin, chitinase and immunity) becoming expressed. Although integration of VLEs into nuclear genomes occurs quite frequently in Saccharomycotina (Frank and Wolfe, 2009; Satwika et al., 2012b; Heneghan et al., 2024), it is unclear whether any of these integrations is transcriptionally active in the nucleus, which probably requires changes to promoter sequences to enable them to be recognized by the nuclear RNA polymerase (Pol II), as well as overcoming mRNA instability caused by the high A+T content of VLEs (Kast et al., 2015). Although both of these scenarios may seem unlikely, one of them must have happened and we think that the second scenario is more plausible than the first.

The sister relationship between Pezizomycotina C-II chitinases and Saccharomycotina VLE chitinases (Figure 4) indicates that although they share a common ancestor, neither of them is directly derived from the other. Otherwise, one of these groups would form a subclade nested inside the other group. Speculatively, it is therefore possible that both groups originated by horizontal transfer from a third, unknown, source such as VLEs in a basal lineage of fungi, which have survived as free VLEs in Saccharomycotina but became integrated into the nuclear genome in an early ancestor of Pezizomycotina. Then, the gene clusters were lost in many Pezizomycotina taxa, but the ones that survived subsequently gained introns, frequently became relocated in the genome and were often duplicated, and some of them became inducible by the presence of other fungi. Their toxins diversified, forming the Tox-P1 to Tox-P9 sequence groups. Further studies will be needed to characterize these Pezizomycotina killer toxins, their potential anticodon nuclease activity and tRNA targets, and their biological roles in interactions among fungi.

## Methods

### Bioinformatics Methods

Chitinases and immunity proteins from the killer VLEs of *Babjeviella inositovora* (pPin1-3), *Debaryomyces robertsiae* (pWR1A), *Kluyveromyces lactis* (pGKL1), and *Millerozyma acaciae* (pPac1-2) were used as queries in TBLASTN searches against Pezizomycotina in the NCBI Whole Genome Shotgun sequences database (Altschul et al., 1994). Any open reading frames having a predicted secretion signal and neighboring a chitinase gene were considered as toxin candidates for further analysis. Additional TBLASTN searches were carried out using the detected Pezizomycotina toxin and immunity candidate proteins as queries. Secretion signals and their cleavage sites were predicted using the TargetP 2.0 webserver (Almagro Armenteros et al., 2019) (https://services.healthtech.dtu.dk/services/TargetP-2.0/). The InterPro database was used to identify the boundaries of the GH18 domains (and Hce2 domains, PF14856) in chitinases. Protein sequences were aligned using Muscle (Edgar, 2004), and phylogenetic trees were constructed by maximum likelihood using Modelfinder in IQ-Tree, with 1000 bootstrap replicates (Minh et al., 2013; Nguyen et al., 2015; Kalyaanamoorthy et al., 2017). Cytoscape visualizations (Franz et al., 2016) were used to display the relationships among toxin candidates, and among immunity protein candidates, including the Saccharomycotina candidates we identified previously (Heneghan et al., 2024). All-versus-all BLASTP searches were used to determine the *E*-value of hits between all toxin pairs, and all immunity protein pairs (Camacho et al., 2009), after which the output was parsed to generate network files for Cytoscape.

### Induction of Candidate Toxins in *Saccharomyces cerevisiae*

Toxin candidates, without their predicted secretion signal sequences, were synthesized by Twist Bioscience with codon optimization for *S. cerevisiae*, and cloned into the integrating plasmid pRG634, which contains a β-estradiol inducible expression system (gift from Robert Gnügge) (Ottoz et al., 2014; Gnugge and Symington, 2020). These plasmids (p141 and p143; Table S7) were transformed into *S. cerevisiae* strain PHY039 (Table S7), a BY4742 derivative, as previously described (Heneghan et al., 2024). Successful transformants were confirmed by PCR. For β-estradiol induction on solid media, *S. cerevisiae* strains containing each toxin gene were streaked on YPD agar ± 2 μM β-estradiol as previously described (Gnugge and Symington, 2020), and photographed after 48 h incubation at 30° C. For induction in liquid media, strains were grown overnight in YPD and then back-diluted to an OD600 of 0.5 in 1 mL of YPD. These were further diluted to an OD600 of 0.1 in 200 μL of YPD ± 0.5 μM β-estradiol in a 96-well plate (each strain, uninduced and induced, was present in duplicate). OD600 was measured every 30 minutes for a 24 h period using a Synergy H1 microplate reader (BioTek).

For galactose induction, the *S. cerevisiae GAL1* promoter was cloned into *Not*I/*Spe*I-digested toxin-pRG634 vectors p141 and p142 (this removes the *LexO-CYC1* promoter; Table S7), and the resulting plasmids (pPGAL1-141 and pPGAL1-142) were transformed into strain PHY039, as described above. For liquid induction, the strains were grown overnight in SC + 2% raffinose, washed twice in PBS, and then each strain was diluted to an OD600 of 0.05 in 200 μL of SC + 2% raffinose or SC + 2% galactose in a 96-well plate. OD600 was measured every 30 minutes for a 24 h period using a Synergy H1 microplate reader (BioTek). For induction on solid media, strains were streaked first to YPD plates and then restreaked to SC + 2% raffinose and grown for 48 h at 30° C. Finally, the strains were restreaked to either SC + 2% raffinose or SC + 2% galactose plates and incubated for 48 h at 30° C.

## Acknowledgements

This work was supported by the European Research Council (789341). We thank Robert Gnügge for plasmids.

## Supplementary Figures and Tables

**Figure S1.**
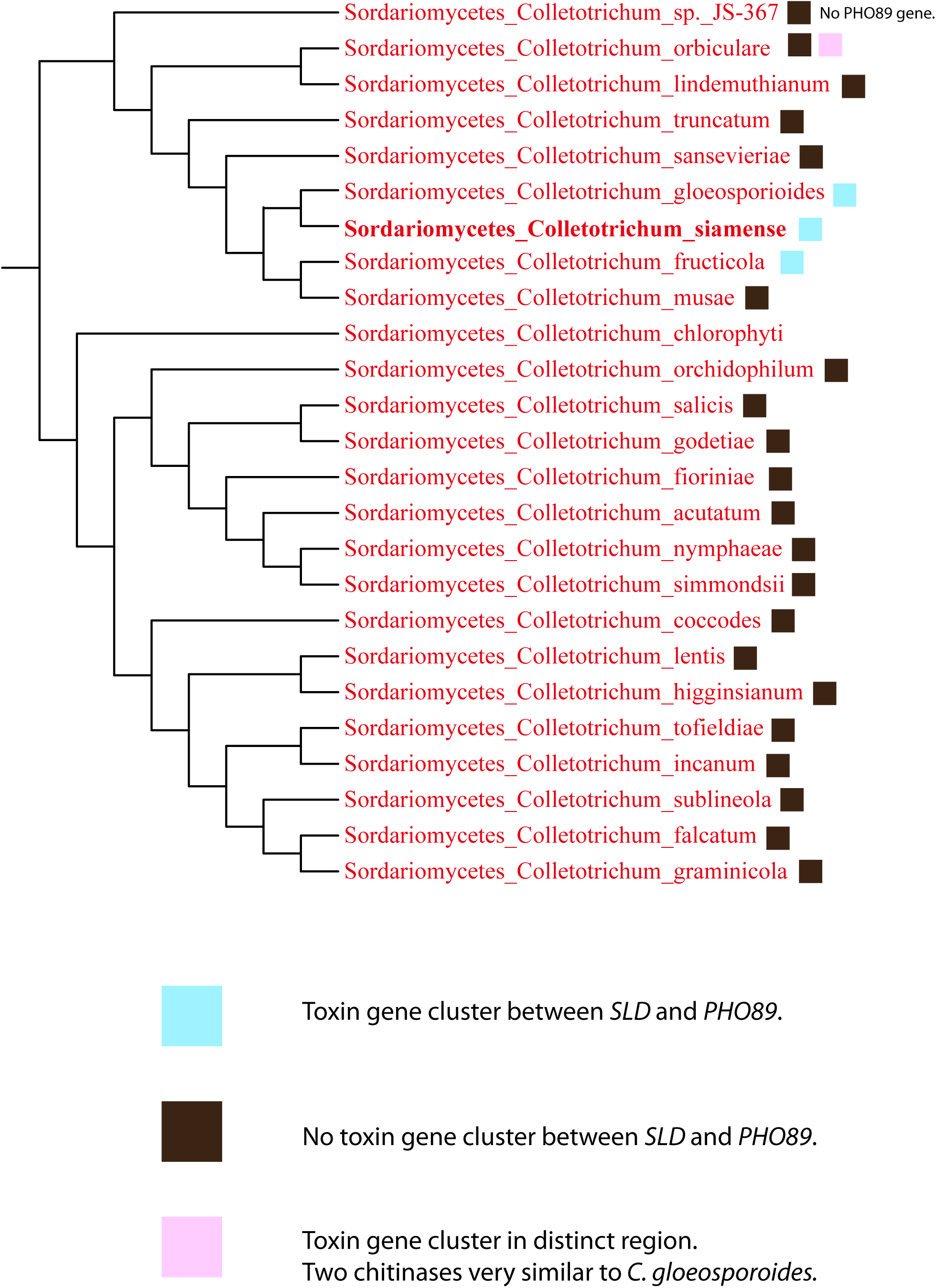
Phylogenetic distribution of a gene cluster in *Colletotrichum* species. The toxin gene cluster in *Colletotrichum siamense* (Figure 1), consisting of genes for a toxin, a chitinase, a LysM-only protein, and a putative immunity protein is also present in two related species: *C. gloeosporoides* and *C. fructicola*. In each of these species, the cluster is between the genes *SLD* (sphingolipid delta-8-desaturase) and *PHO89* (phosphate transporter). However, none of the other *Colletotrichum* species have a cluster at the syntenic location. The topology of the phylogenetic tree is based on the fungal phylogenomic tree constructed by Shen et al. (2020). *C. siamense* was not included in Shen et al’s tree because its genome was sequenced too recently, so it is shown at an estimated position based on the fact that it is a member of the *C. gloeosporoides* species complex (Rehner et al., 2023).

**Figure S2.**
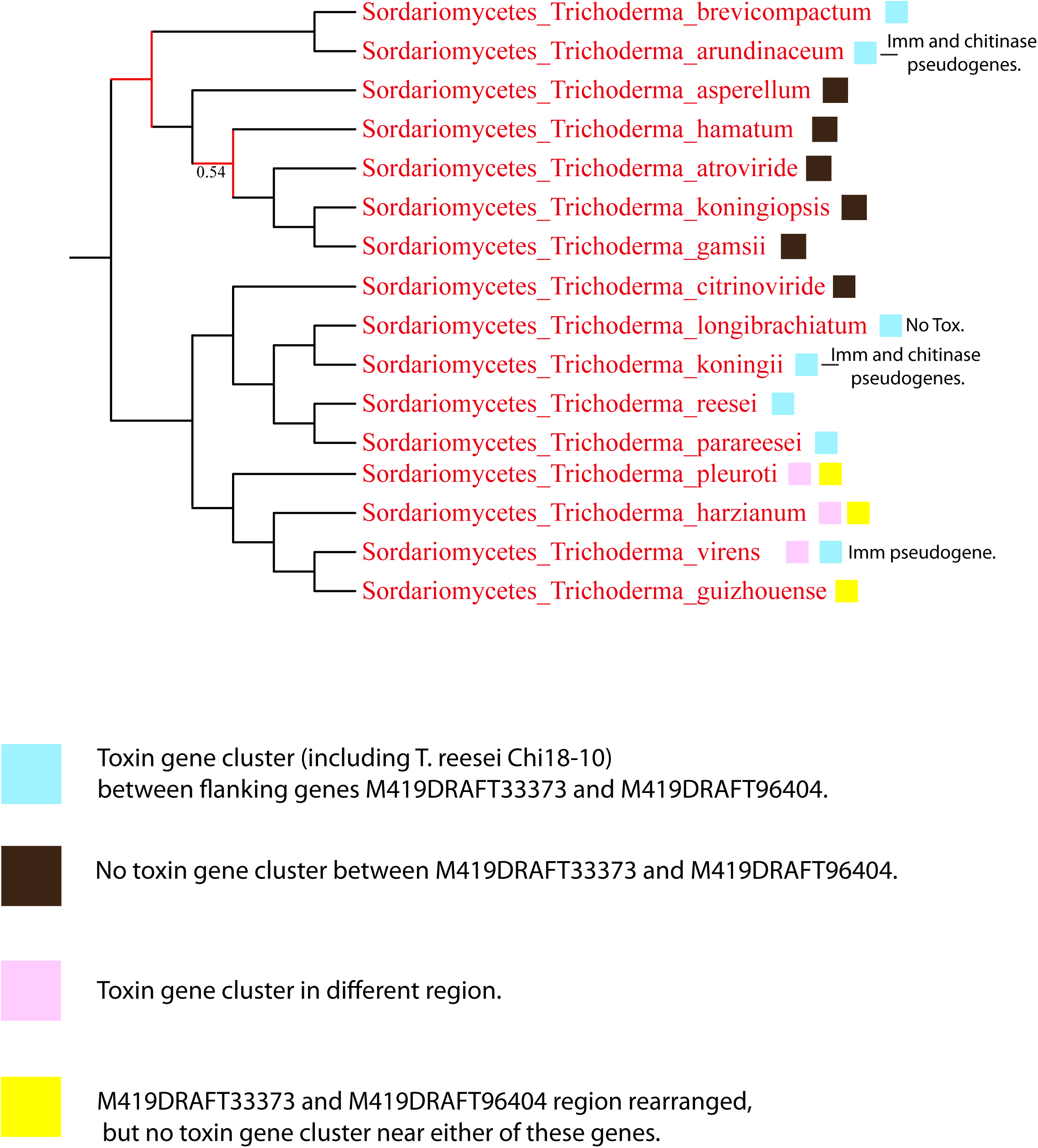
Phylogenetic distribution of a gene cluster in *Trichoderma* species. We tracked the evolutionary origin of the toxin gene cluster beside chitinase *Chi18-10* in relatives of *Trichoderma reesei*. The genes flanking the cluster have the systematic names *M419DRAFT33373* (serine/threonine kinase) and *M419DRAFT96404* (mitochondrial ornithine transporter) in the *T. reesei* genome sequence annotation. This cluster appears to have been present in the common ancestor of the genus *Trichoderma*, but it has been lost in some species and moved to a different genomic location in others. The species phylogeny is from Shen et al. (2020).

**Figure S3.**
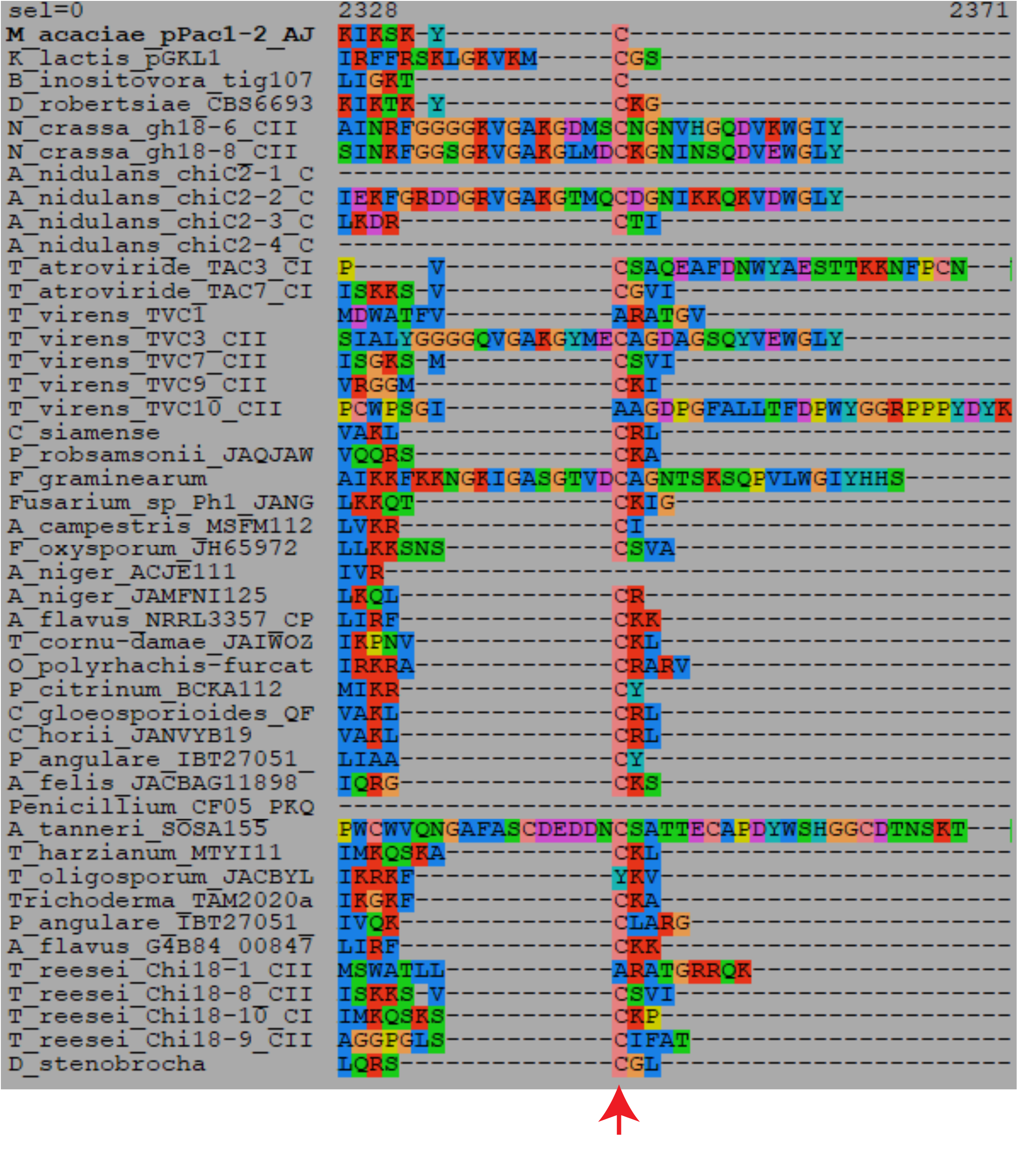
Presence of a Cys residue near the C-terminus of Pezizomycotina chitinases. The C-terminal region is shown of an alignment of Pezizomycotina chitinases (from Figure 1), as well as the chitinase (killer toxin α/β subunit precursor) from *Kluyveromyces lactis* pGKL1, *Debaryomyces robertsiae* pWR1A, *Millerozyma acaciae* pPacl-2, and *Babjeviella inositovora* pPinl-3. The red arrow indicates the cysteine residue.

**Figure S4.**
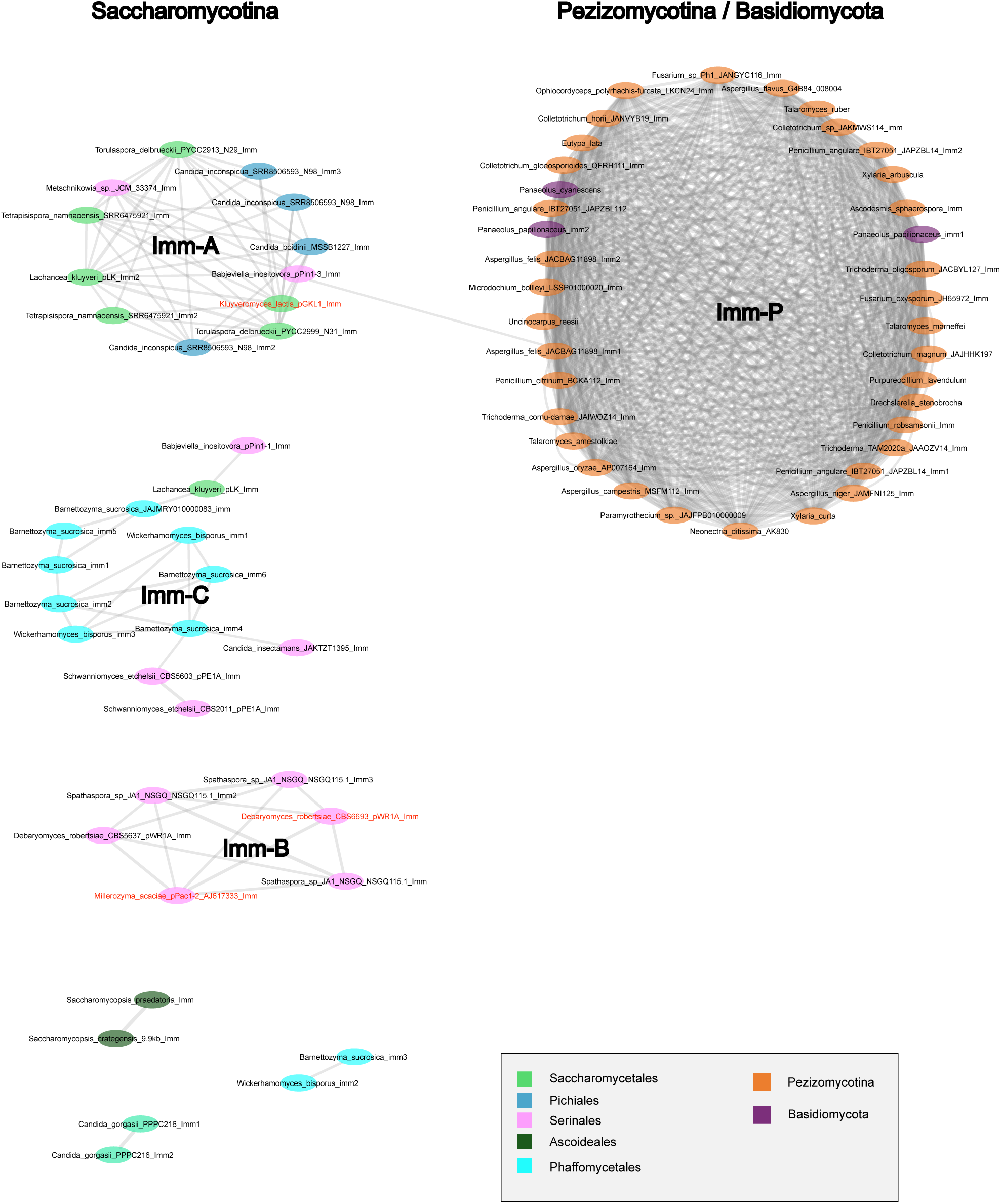
Network of putative immunity proteins from Saccharomycotina and Pezizomycotina. Each node represents an immunity protein. Proteins are colored by taxonomic group as shown in the key. Two nodes are connected by an edge if the BLASTP hit between them has a significance of *E* < 1e-13. The Pezizomycotina proteins fall into a single large cluster, labeled Imm-P. The previously identified Saccharomycotina immunity protein sequence groups Imm-A to Imm-C are labeled (Heneghan et al., 2024). At this BLASTP cutoff, there is only a single hit between a Saccharomycotina protein and a Pezizomycotina protein.

**Table S1. Details of Pezizomycotina chitinase genes and gene cluster locations.**

**Table S2. Locations of candidate immunity genes in Pezizomycotina genome assemblies, not located in clusters.**

**Table S3. Details of toxin candidates found in Pezizomycotina.**

**Table S4. Details of candidate Immunity genes found in Pezizomycotina.**

**Table S5. Details of chitinase genes in Basidiomycota gene clusters.**

**Table S6. Details of toxin candidates found in Basidiomycota.**

**Table S7. Strains and plasmids used in this study.**

